# Evolution engineering of methylotrophic E. coli enables faster growth than native methylotrophs

**DOI:** 10.1101/2024.04.18.589993

**Authors:** Liang-Yu Nieh, Frederic Y.-H. Chen, Hsin-Wei Jung, Kuan-Yu Su, Chao-Yin Tsuei, Chun-Ting Lin, Yue-Qi Lee, James C. Liao

## Abstract

As methanol can be derived from either CO_2_ or methane, methanol economy may play a role in combating climate change. In this scenario, rapid utilization of methanol by an industrial microorganism is the first and crucial step for efficient utilization of the C1 feedstock chemical. Here, we report the development of a methylotrophic E. coli strain (SM6) with a doubling time of 3.5 hours, outpacing that of common native methylotrophs. We accomplish this using evolution engineering with dynamic copy number variation (CNV). We developed a bacterial artificial chromosome (BAC) with dynamic CNV to facilitate overcoming the formaldehyde-induced DNA-protein cross-linking (DPC) problem in the evolution process. The growth rate of the organism in methanol minimal medium improved significantly after it acquired a loss-of-function mutation in *mutS*. We tracked the genome variations of 72 cultures along the evolution process by next-generation sequencing, and identified the metabolic features of the fast-growing strain. This study illustrates the potential of dynamic CNV as an evolution tool and synthetic methylotrophs as a platform for sustainable biotechnological applications.

## Introduction

Methanol is the most reduced C1 compound that exists in the liquid form in most relevant conditions ^1,2^. This property by-passes many difficulties in transportation and storage of C1 compounds. As such, use of methanol as a starting point for chemical production has been proposed as a potential route to reduce greenhouse gas accumulation, since methanol can be derived from CO_2_ or methane. In this regard, developing industrial microorganisms that can use methanol for growth and production has been a goal for many groups ^3–12^.

Rapid growth of microorganisms in methanol minimal medium is an essential step and the first requirement in microbial conversion of methanol. However, current synthetic methylotrophic strains evolved from non-C1-utilizing microbes still grow much slower than typical industrial strains^5,7,9,13^. Laboratory evolution, a widely adopted approach in engineering microorganisms, is particularly suitable for improving growth rate since it can enrich fitter organisms^4–6,8,13,14^. Methanol-utilizing synthetic methylotrophs face a particular problem, formaldehyde-induced DNA-protein crosslinking (DPC) caused by imbalance between methanol oxidation that leads to formaldehyde production and formaldehyde consumption^5^. These synthetic organisms may accumulate and release formaldehyde during later growth phases, ultimately leading to self-destruction and poisoning surrounding cells. This scenario impedes beneficial mutations from accumulating during the evolution in the presence of methanol. This may be one of the underlying reasons for the difficulty in engineering or evolving a fast-growing synthetic methylotroph to date.

Formaldehyde is an essential intermediate in methanol utilization, but its production and consumption are difficult to balance at all physiological stages. Native methanol-utilizing organisms likely have evolved sophisticated regulatory mechanisms to balance the formaldehyde production and consumption. One such mechanism is the glutathione-dependent formaldehyde oxidation pathway that converts formaldehyde to formic acid and serves as a surge release^15^. However, this pathway reduces the biomass yield of the methanol-grown strain and is not desirable for our purpose here. In fact, this protective pathway was deleted in multiple reported synthetic methylotrophs^4–6,8,10^.

Without the protective formaldehyde oxidation pathway, this intermediate is likely to accumulate, as we push for fast methanol conversion. Once formaldehyde accumulates, it causes the DPC problem that kills the producing cells. Formaldehyde can also diffuse in the medium and cause DPC in other cells. Thus, even if the cell is engineered to utilize methanol for fast growth under a particular condition, formaldehyde accumulation may occur at a later stage and cause DPC to kill itself and others. Consequently, cells with beneficial mutations are difficult to enrich or accumulate. Moreover, cells in liquid culture or colonies on solid medium cannot be re-cultured until a long lag phase has passed^5^. Since all DNA and proteins are susceptible to DPC, formaldehyde-resistant cells are difficult to emerge. The DPC problem is particularly severe in high methanol concentrations.

In this work, we developed a dynamic CNV scheme to facilitate de-bottlenecking the laboratory evolution process in high methanol concentrations. After de-bottlenecking, the strain was able to evolve and developed a *mutS* allele which accelerated mutation. Eventually a single colony was isolated (SM6) with a doubling time (T_d_) of 3.5 hours, demonstrating faster growth than native methylotroph, *Methylorubrum extorquens* AM1 (T_d_∼4hr)^16^ and *Bacillus methanolicus* (T_d_ ∼ 5hr at 37°C.). Finally, we characterized the metabolic features of SM6 to find the reason underlying its fast growth.

## Results

### Dynamic Copy Number Variation Introduced by ddp-BAC

To address the DPC issue that hinders long-term evolution, we developed a dynamic CNV scheme to tune the expression levels of formaldehyde consumption enzymes. This scheme is based on our previous discovery ^5^ of a tandem repeated 7kb region in the strain (BB1) (see Supplementary Fig, 1 for BB1 7kb tandem repeats) which spontaneously emerged and co-evolved with our synthetic methylotrophic SM1 strain that can utilize methanol as the sole carbon source for growth (see Supplementary Fig. 2 for ^13^C labeling data). We cloned this operon onto a bacterial artificial chromosome (BAC) with a *gfp* expression cassette. Surprisingly, we found that the presence of the *ddp* operon in the ddp-BAC spontaneously developed tandem repeats without selection pressure, forming a circular concatemer of the whole ddp-BAC (Fig. 1 abc). Furthermore, we found that the concatemer and CNV disappeared in a *ΔrecA* background (Fig. 1bc), indicating a *recA*-dependent process in CNV. This *recA*-dependent process can occur within a cell, as demonstrated by time-lapse light microscopy (Fig. 2a), Using flow cytometry, we found that the ddp-BAC::*gfp* exhibited a bimodal distribution (Fig. 2b) with a wide-range of fluorescence intensity, which presumably corresponds to copy number distribution.

**Figure 1.**
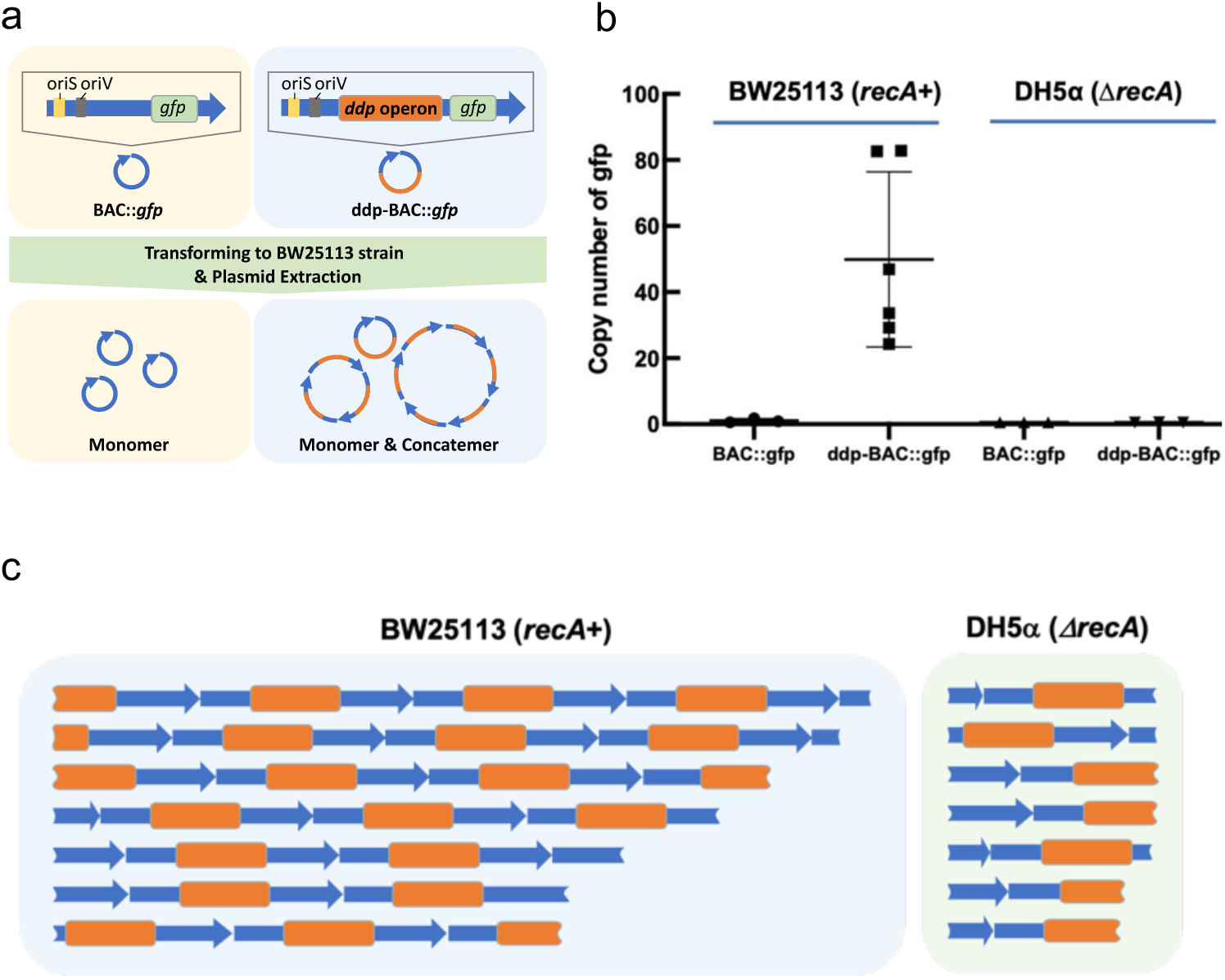
Copy number and tandem repeat pattern of BAC. (a) Cartoon scheme of ddp-BAC and BAC. (b) Copy number of *gfp* measured by droplet digital PCR (ddPCR). Only BAC with a *ddp* operon in a *recA*^+^ environment exhibits a high copy number. (ddp-BAC: n=6; others: n=3, biological repeats) (c) Nanopore sequencing of ddp-BAC. The pattern shown are raw, intact nanopore reads that were mapped to BAC. The orange label represents the *ddp* operon, while the blue arrows represent the BAC backbone with *gfp*. Only ddp-BAC in *recA*^+^ host showed tandem repeats with a distribution of copy numbers.

**Figure 2.**
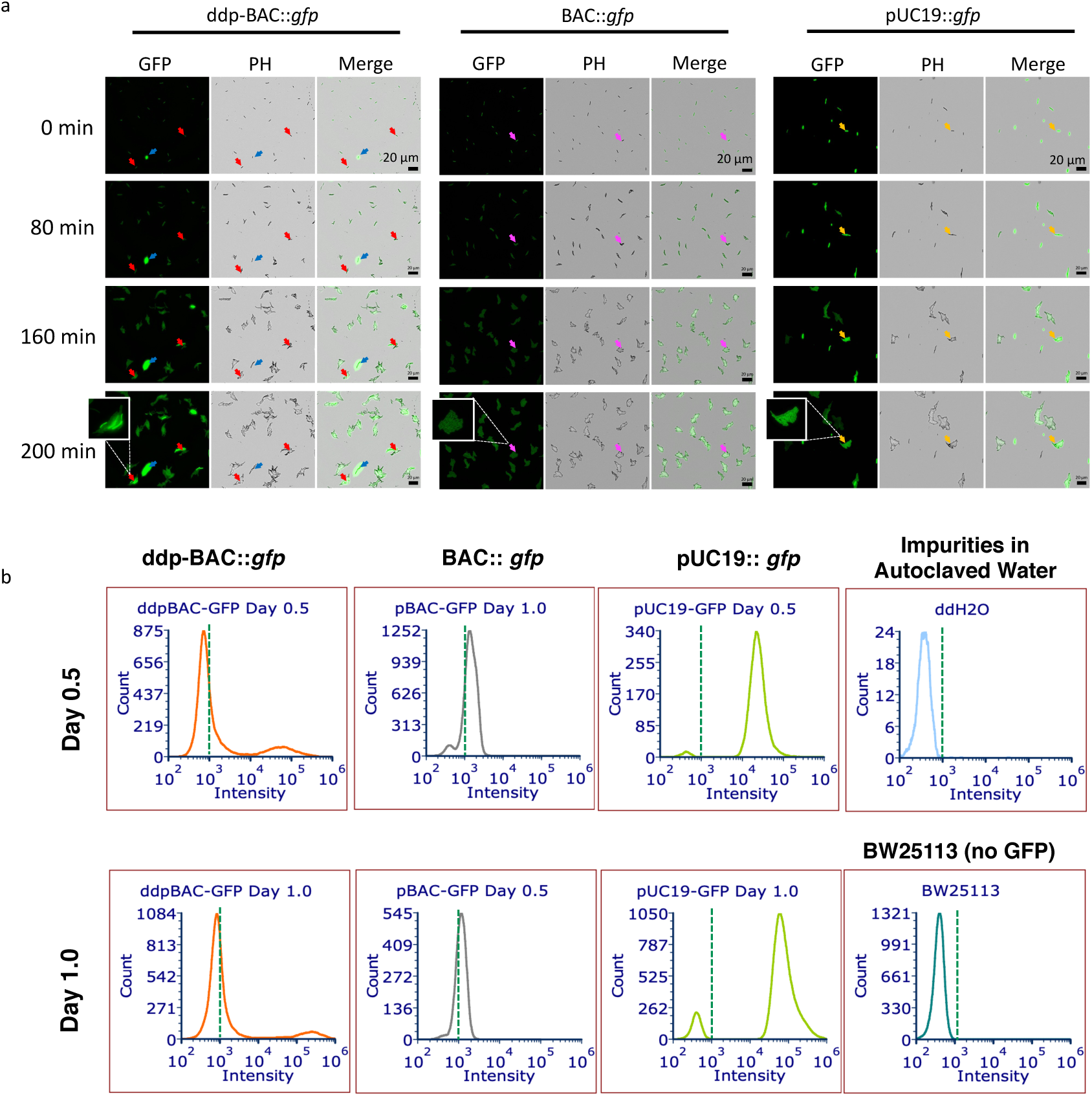
Copy number distribution of ddp-BAC. (a) Time-lapse live cell imaging of GFP fluorescence. Colonies from ddp-BAC::*gfp* showed varied fluorescence levels both among different colonies and within the same colony. The red and blue arrows indicate low and high fluorescence, respectively. In contrast, colonies from strains BAC::*gfp* (magenta arrows) or the high-copy pUC19::*gfp* (orange arrows) demonstrated uniform fluorescence intensities. (b) Flow cytometry analysis of BW25113 containing various versions of BAC or pUC19. Autoclaved water and WT BW25113 were used to determine the GFP fluorescence threshold. Results showed that only ddp-BAC::*gfp* exhibited a bimodal distribution.

The mechanisms underlying the dynamic CNV is currently under investigation and will be reported elsewhere. Nevertheless, this wide distribution of copy numbers within a population and the dynamic tuning of copy number by RecA could potentially be useful in laboratory evolution. We thus apply this property in improving the growth rate of synthetic methylotrophic *E. coli*.

### Use of ddp-BAC to de-bottleneck evolution

While the synthetic methylotrophic *E. coli* SM1 that we previously developed^5^ has alleviated the DPC problem, its growth rate (T_d_= 8.5 hr) is still far below that of the native methylotroph or *E. coli* growth rate in glucose. This slow growth was presumably due to the residual DPC issue. Indeed, the residual DPC issue was amplified when the strain was grown in high methanol concentrations (Fig. 3a). We initially evolved the strain for 81 passes, but the improvement of growth rate was marginal and genomic changes were minor (Fig. 3b). Therefore, we decided to use the ddp-BAC as a tool to tune the copy number of the formaldehyde consumption enzymes.

**Figure 3.**
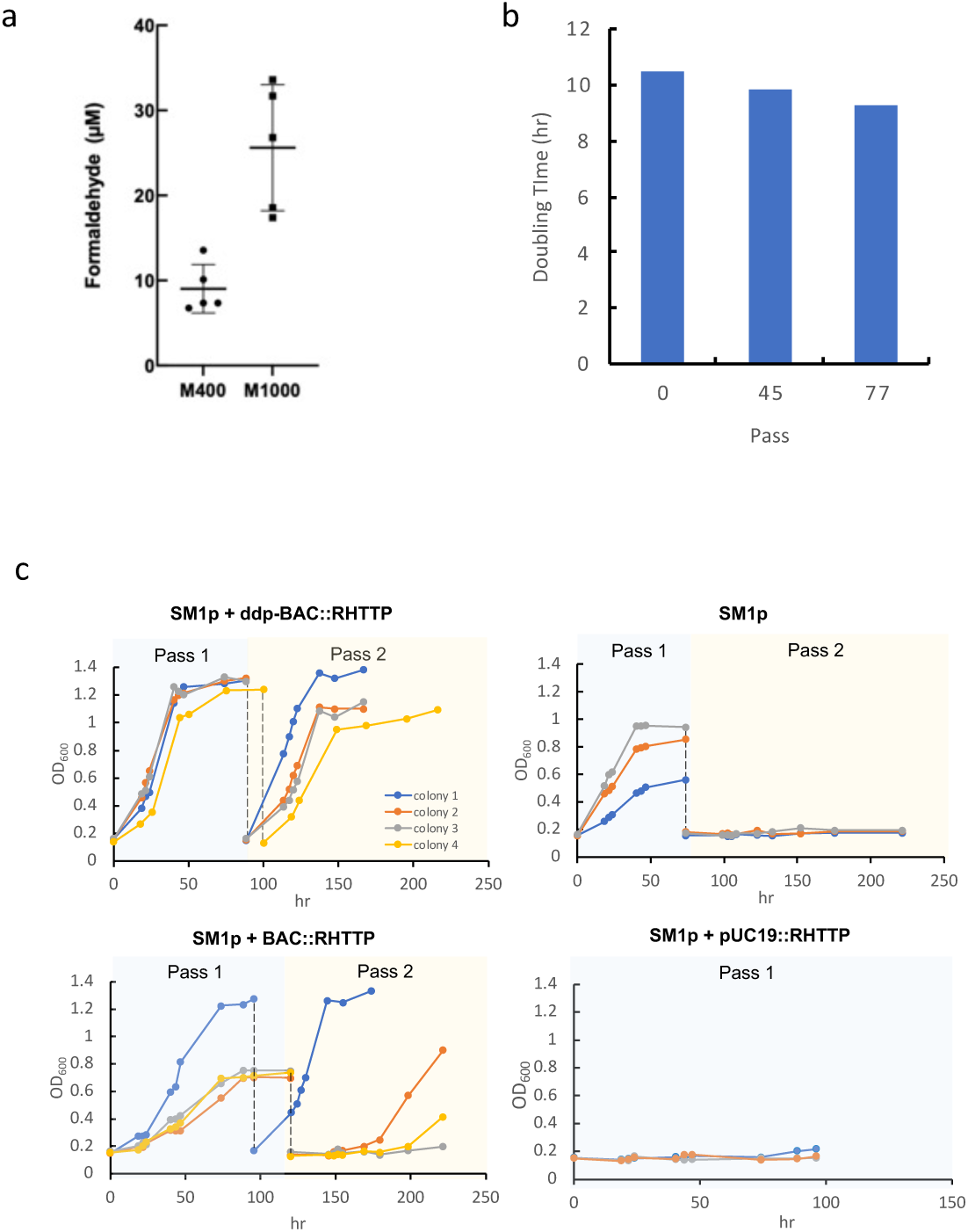
De-bottlenecking evolution using ddp-BAC::RHTTP. (a) Formaldehyde concentration in SM1 cultures OD 1.0 in low (400mM, M400) and high (1000mM, M1000) methanol concentrations (n = 3 for each group, biological repeats; error bars= S.D.). (b) Doubling time of the SM1 strain evolved up to pass 77 showed minor improvement. (c) SM1 with various plasmids harboring RHTTP grown and passed in 1000mM methanol. Each strain was tested with at least 3 colonies from an LB plate.

We cloned the genes for formaldehyde utilization in the ribulose monophosphate (RuMP) cycle—*rpe*, *hps*, *tkt*, *tal*, and *phi* (collectively RHTTP) on the BAC system with or without ddp and transformed into strain SM1p, which is a descendent of SM1 after 81 passes (equivalent to about 284 generations) of laboratory evolution in 400 mM of methanol minimal medium. Additionally, this strain grew poorly in high methanol medium (1000 mM) and could not continue to grow in the second pass (Fig. 3c), suggesting a severe DPC problem in high methanol medium that killed the cells.

In contrast, the strain containing ddp-BAC::RHTTP system (pLY147) grew faster in 1000 mM of methanol, and can continue to grow without a lag phase in the subsequent passes (Fig. 3c). This may suggest that the ddp-BAC::RHTTP system effectively resolved the DPC issue in the high methanol medium. As a comparison, the strain containing BAC::RHTTP (pLY143, without the ddp operon) grew slightly slower and exhibited a lag phase in the second pass (Fig. 3c) The beneficial effect of ddp-BAC::RHTTP is presumably attributed to the high copy number of RHTTP on the ddp-BAC system. To test if a high copy number plasmid (pUC19) can achieve the same effect as the ddp-BAC system, we also expressed RHTTP using pUC19. Interestingly, the pUC19::RHTTP system did not allow the strain to grow in 1000 mM of methanol (Fig. 3c), presumably because of large protein burden.

Thus, it appears that the bimodal distribution of copy numbers and the *recA*-dependent tuning of copy number allows the strain to find an optimal expression level to by-pass the DPC problem in high methanol concentrations.

### Evolution to develop a fast-growing methylotrophic *E. coli* strain

We then used the ddp-BAC::RHTTP system to continue the evolution in 1000 mM methanol for 18 passes, and observed the improvement of growth rate (Fig. 4a). Then, we raised the methanol concentration to 1200 mM to increase the selection pressure. Again, the growth rate in high methanol concentration continued to improve. At passage 132, we sequenced the strain and found that the *recA* gene on the chromosome was interrupted by an insertion sequence (IS2). As a result of the *recA* inactivation, the copy number of the RHTTP gene on the ddp-BAC decreased to 1. This result once again confirmed the *recA*-dependent CNV of ddp-BAC, and suggested that during the evolution, the initial high copy number of RHTTP (Fig. 4a) allowed the strain to avoid DPC and continued to evolve, even though the high copy of RHTTP may introduce a protein burden. During the subsequent evolution process, the cell managed to accumulate other mutations and did not need the high copy RHTTP. Thus, an IS2 insertion of *recA* was enriched and dominated the culture.

**Figure 4.**
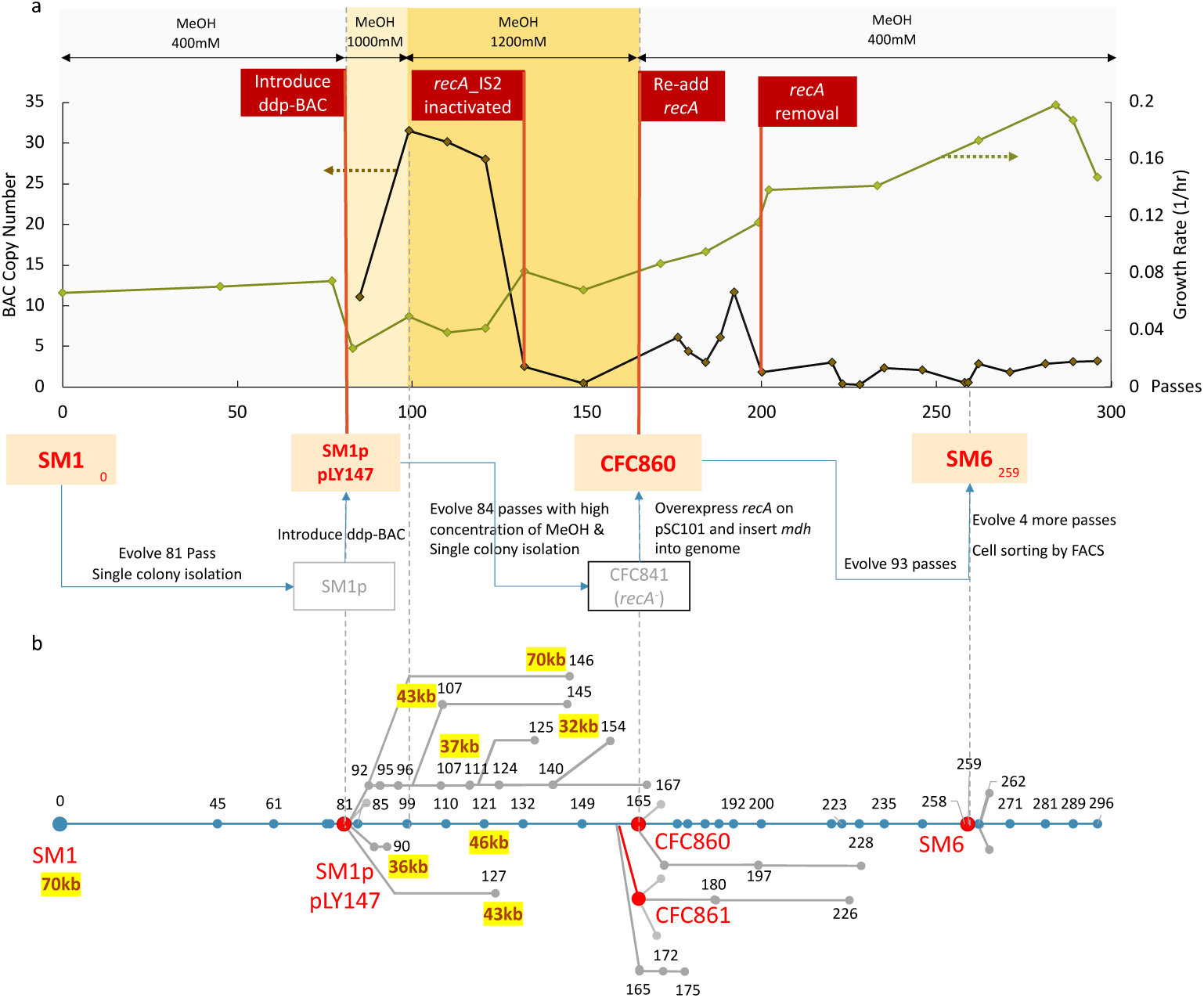
The evolution process towards SM6. (a) Copy number of ddp-BAC and growth rate change along the evolution line towards SM6. The copy number of ddp-BAC was derived from Illumina sequencing. Note that the copy number significantly dropped when *recA* was interrupted or removed. (Each “pass” is defined as a subculturing step, growing from OD 0.05∼0.1 to OD1) (b) The overall evolution lineage graph with branching evolution lines. The numbers depict the passage number of the strain. All dots represent samples that were sequenced by NGS. The yellow highlighted numbers depict the unit size of the IS5-flanked tandem repeats derived from the original 70kb tandem repeat.

At this point, the strain presumably has found ways to avoid the DPC problem. We then focused on increasing methanol consumption by introducing another copy of *mdh* to the chromosome, and returned to the evolution process in low methanol concentration (400mM). At 259 passes, we finally identified a culture which has a doubling time of less than 4 hours. From this culture, we isolated a single strain, named SM6, which has a T_d_ of 3.5 hr (growth rate = 0.2 hr^-1^) (Fig. 4a), which is faster than model methylotrophs *Methylorubrum extorquens* AM1 (T_d_∼4hr) and *Bacillus methanolicus* (T_d_∼5hr).

### Tuning of copy number by *recA* inactivation

To further understand the genomic variations during the evolution process, we sequenced a total of 72 cultures using Illumina Mini-seq along the main evolution line and in the branched evolution lines (Figs. 4 & 5).

**Figure 5.**
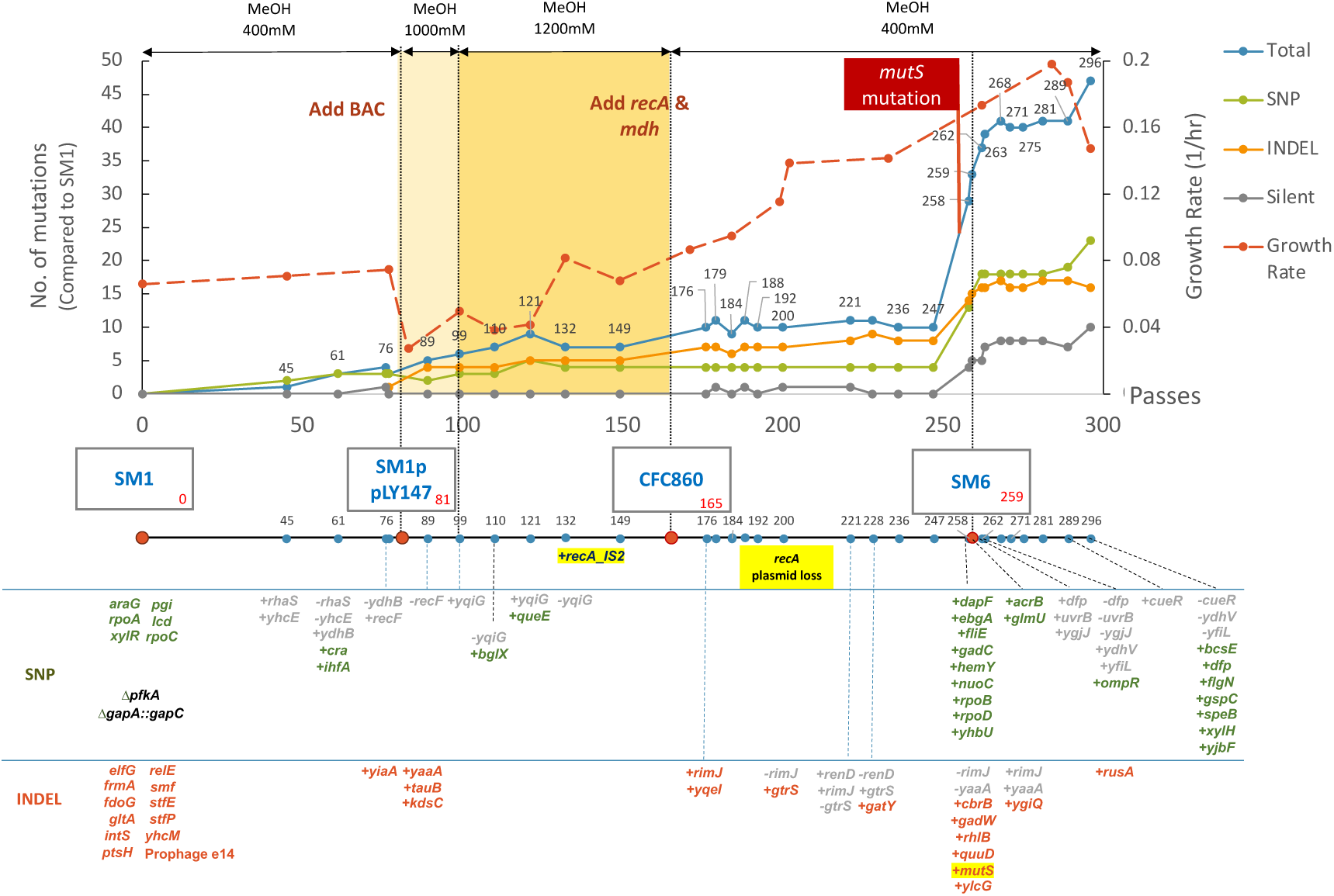
Mutation timeline during the evolution process. Only changes greater than 25% variant frequency are shown. The genomic changes in SM1 are relative to its parental strain BW25113. The “+” & “-” signs represent mutations appeared or disappeared along the evolution line, relative to the starting strain SM1. The IS2-inserted *mutS* inactivation coincided with the increase of mutation events. All genes that were not retained along the evolution process are labeled in grey.

As expected, the copy number of ddp-BAC::RHTTP increased after its introduction, and reached 30 copies within 20 passes in high methanol concentration, then rapidly decreased to 1, which coincided with a *recA* insertion by IS2 (Fig. 4). Presumably, during this period of time, the cell accumulated sufficient beneficial genomic variations, and do not need the high copy RHTTP anymore. Thus, a *recA* inactivation was enriched to reduce the copy number to 1 and avoid protein burden.

To test this hypothesis, we introduced a *recA*-containing plasmid pKY29 at 165 passes (Fig. 4a). The introduction of the active *recA* gene indeed increased the copy number of RHTTP to 12 in 30 passes. However, within 5 additional passes, the *recA*-containing plasmid was removed and the copy number of RHTTP was reduced to 2 (Fig. 4a). Thus, it is evident that the cell used RecA and ddp-BAC to tune the copy number of RHTTP that fits the genetic background. Since genomic changes continued to emerge along the evolution line while the copy number of ddp-BAC decreased (Fig. 4a, Fig. 5, Supplementary Fig. 3), it was possible that the ddp-BAC provided a favorable background for the cell to accumulate beneficial mutations while decreasing its copy number for better fitness.

### Genomic changes during the evolution process

Next, we examine the genomic landscape during the evolution process. The parental strain SM1 contained 5 copies of tandem repeats each spanning 70kb flanked by IS5 inserted next to *yggE* and *yghO*. Interestingly, the IS5-mediated tandem repeat unit was shrunk from 70kb to 46kb at pass 121 without changing the copy number (Supplementary Fig. 3). One of the IS5 boundaries next to *yghO* gene was present in the ancestral BW25113 strain and the parental SM1 strain, and remained unchanged. The other IS5 boundary next to *yggE* was transposed from its original position to *endA*. The shrinking of the 70kb tandem repeats was also observed in other branches of evolution, resulting in tandem repeat units with size ranging from 32kb to 46kb (Fig. 4b, Supplementary Fig. 3). Curiously, these events all occurred after introducing the ddp-BAC::RHTTP system.

Another key genomic change during the main evolution line was the emergence of the IS2 insertion into *mutS* which occurred at 258 pass (Fig.5). MutS is responsible for excision mismatch repair, thus its inactivation caused an elevated mutation rate^17^. After this event, the number of SNP and INDEL rapidly increased, and the growth rate improved (Fig. 5). We then isolated the fastest growing strain from 258 pass, which is SM6. Using the rifampicin assay, we confirmed that the mutation rate of SM6 is approximately 80-fold higher than that of SM1 and the BW25113 wild-type strain (Table 1). Further evolution of SM6 continued to accumulate mutations, but the growth rate eventually decreased.

**Table 1.** A rifampicin-based mutation assay reveals that SM6 showed an 80-fold higher mutation rate compared to that of the wild-type and SM1 strains.

| Strain | Cell number grown on rifampicin | Total cells (CFU/mL) | Fold change of mutation rate |
| --- | --- | --- | --- |
| BW25113 | 120 | $3.5 \times 10^9$ | 1 |
| SM1 | 11 | $0.2 \times 10^9$ | 1.6 |
| SM6<br>(*mutS* mutation) | >1000 | $0.4 \times 10^9$ | >80 |

### Characterization of SM6

To confirm that SM6 had solved the DPC issue, we compared the extracellular formaldehyde concentration of SM1 and SM6 at different growth stages (Fig. 6ab). Compared to SM1, the concentration of formaldehyde in SM6 cultures consistently remained low, approximately around 20 µM. This suggests a well-balanced flux between formaldehyde production and consumption, regardless of the growth stage. The absence of DPC complexes in different growth stages of SM6 were then confirmed by transmission-electron microscopy (TEM) (Figure 6b).

**Figure 6.**
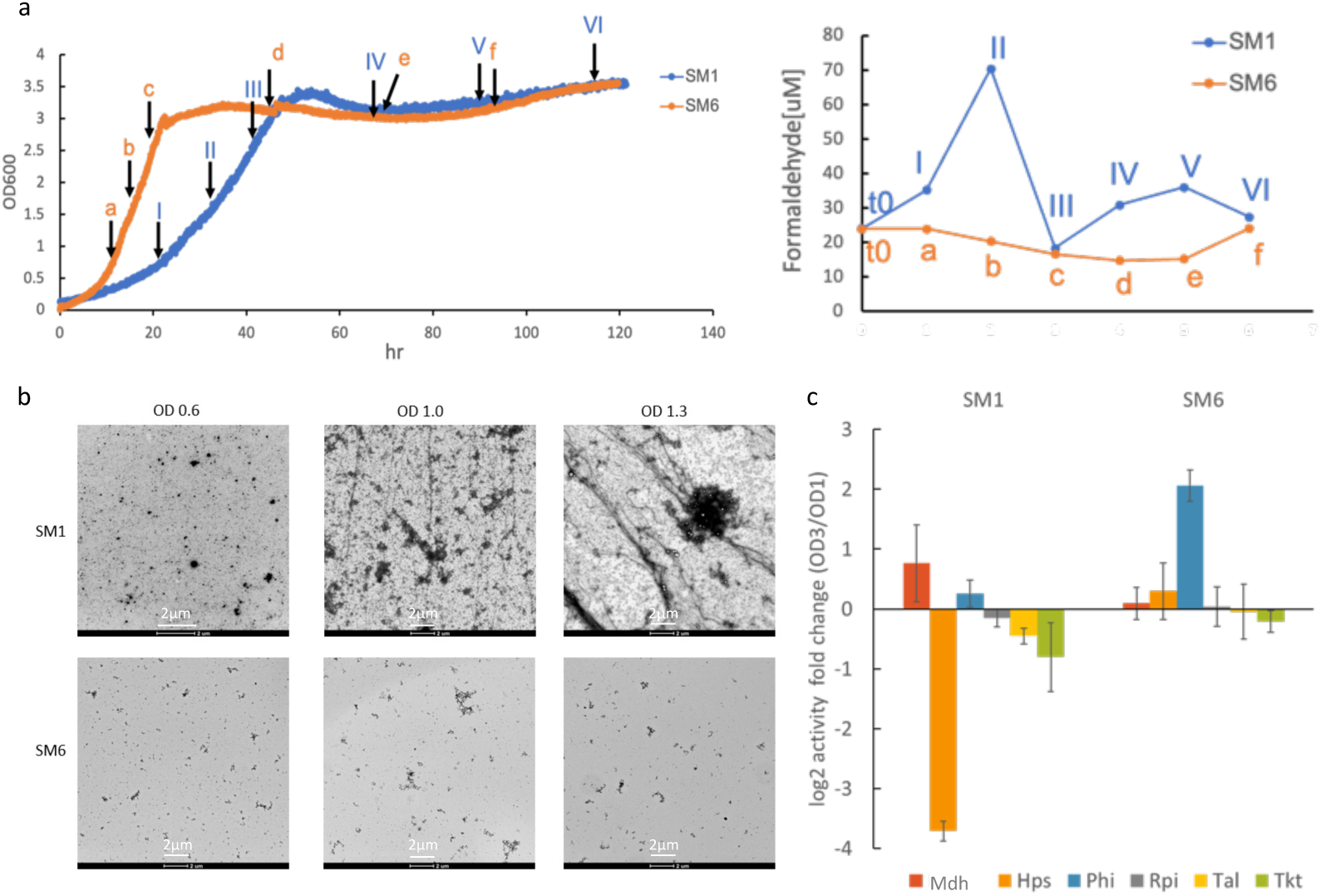
Characterization of DPC in SM6. (a) Time courses of SM1 and SM6 growth. Samples were collected at time points I to VI for SM1, and a to f for SM6, respectively, for formaldehyde measurement using LC-MS/MS. (b) Transmission electron microscopy (TEM) images of DPC products extracted from both SM1 and SM6 at different growth stages by negative staining. (c) Enzymatic assay of Mdh and enzymes in the RuMP cycle in SM1 and SM6. Mdh activity was measured using the Nash assay. The remaining enzymes were measured using coupled enzyme assays with a NADPH readout at 340nm (n=3, biological repeats; error bars= S.D.)

We also assayed the activities of enzymes related to formaldehyde production and consumption, including methanol dehydrogenase (Mdh), hexulose-6-phosphate synthase (Hps), 6-phospho-3-hexuloisomerase (Phi), transketolase (Tkt), transaldolase (Tal) and ribose-5-phosphate isomerase (Rpi) (Fig. 6c). The results indicated that in SM6, Hps, Phi, exhibited a significant increase as the strains entered the stationary phase. This increase in formaldehyde consuming enzyme expression in the stationary phase avoided the DPC problem. In contrast, the SM1 strain showed a significant decrease in Hps, Tal, Tkt, while Mdh increased during the stationary phase, causing the imbalance of formaldehyde production and consumption. Although the DPC issue in SM1 was already improved from its parental strain, the residual DPC may still impedes growth.

We also characterized the proteome SM6. Strikingly, the ribosome abundance increased 50% compared to SM1 (Fig. 7a). The increase in ribosome abundance may be a key reason underlying the fast growth of SM6. In addition, the enzymes in amino acid pathways were more abundant in SM6 than in SM1 (Fig. 7b), which may also contribute to the fast growth of SM6. Comparing the proteomes of SM6 and the wild-type parental BW25113, we found that the RuMP cycle enzymes were increased as expected. However, the AceEF, PflB, GltA decreased while PoxB, Acs and GlcB increased (Fig. 7c). These changes suggest that SM6 derives 2-carbon compounds not from pyruvate dehydrogenase (coded by *aceEF*) or pyruvate-formate lyase (coded by *pflB*) but from pyruvate oxidase (coded by *poxB*) and acetate utilization (coded by *acs*, *glcB*).

**Figure 7.**
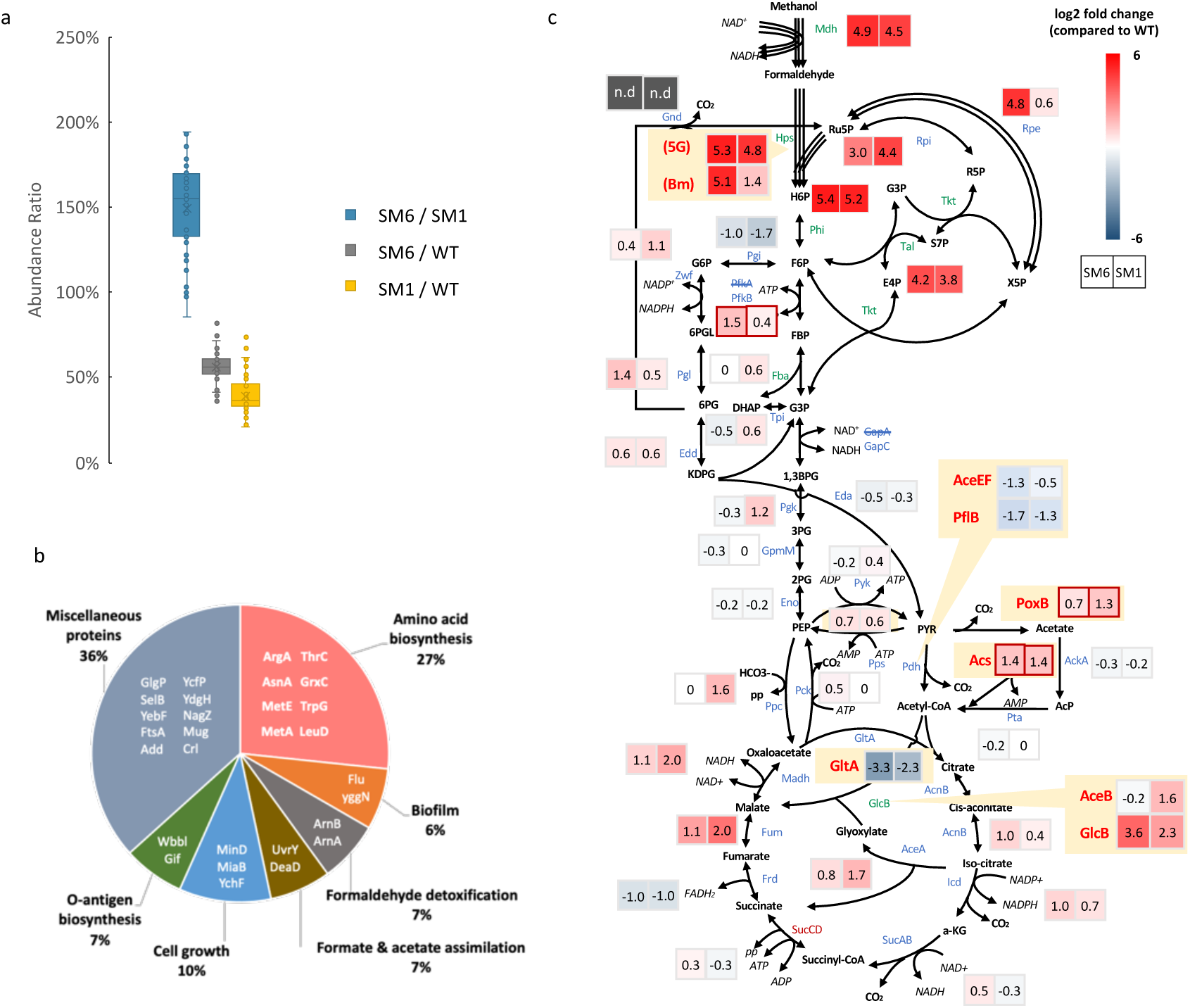
Proteomics analysis of SM6 and SM1. (a) Abundance ratio of the 44 *E. coli* ribosomes in SM6, SM1 and wild-type parental strains. (error bar = S.D.) (b) Functional enrichment analysis identified biological processes in the top 30 up-regulated proteins by comparing SM6 to SM1 grown on 400 mM methanol until OD_600_ reached 1. (c) Metabolic protein expression profiles of SM6 (left box) and SM1 (right box) relative to the wild-type parental strain. Gray blocks indicate not detected value (n.d) due to gene knockout. Hps were cloned from either *Methylomicrobium buryatense* 5GB1S (5G) or *Bacillus methanolicus* (Bm).

The final SM6 strain contains a ddp-BAC::RHTTP, but because of the inactive RecA, the ddp-BAC::RHTTP exhibited only 2 copies. To probe the effect of this system on SM6 growth, we rid of ddp-BAC::RHTTP from SM6 (renamed SM6n) and found that the strain grew slower (T_d_=7 hr) (Fig. 8a). This result validated the contribution of ddp-BAC::RHTTP in SM6.

**Figure 8.**
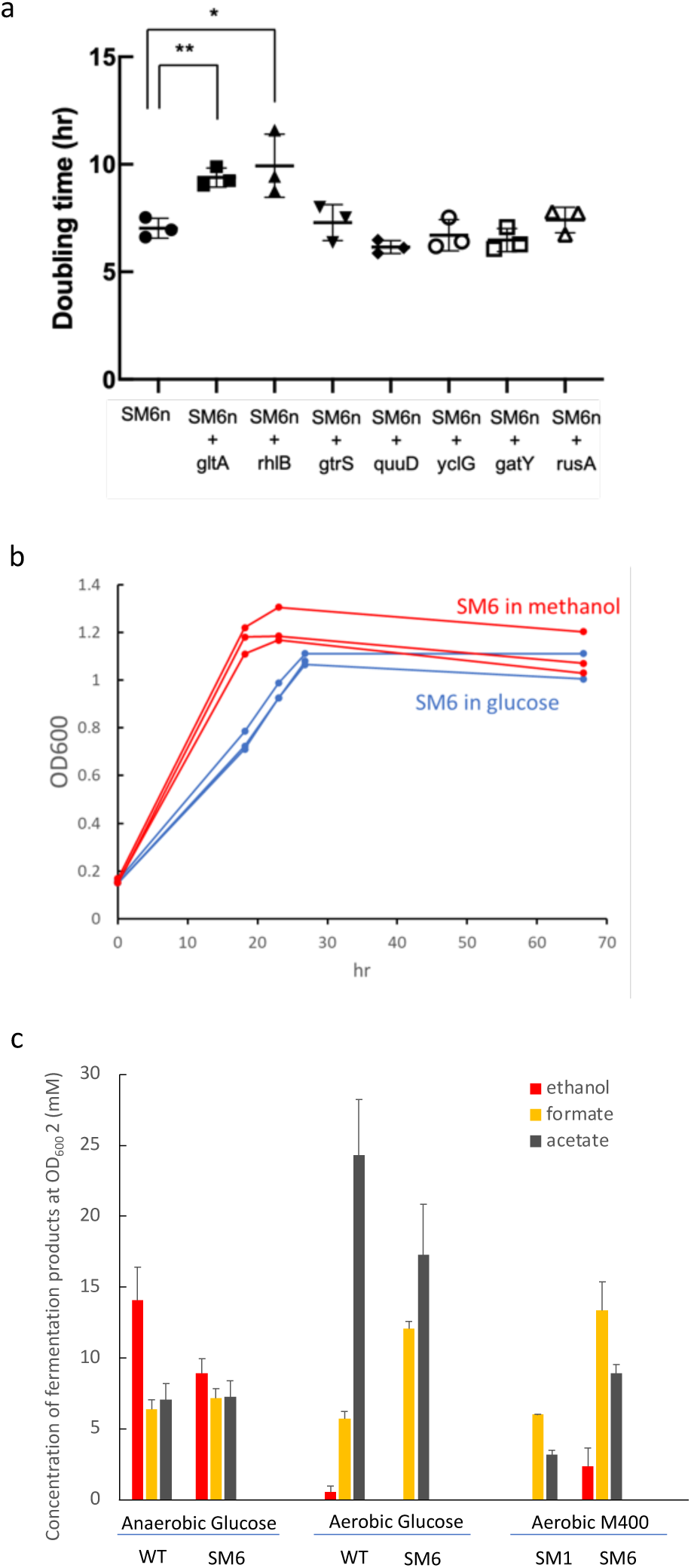
SM6 characteristics. (a) Complementation of mutated genes with BAC harboring a wild-type copy of the indicated genes. Growth rate of SM6n with and without BAC complementation were measured (n = 3 for each group, biological repeats; error bars= S.D.; significance was tested by 2-side Welsh-test). (b) The growth rate SM6 in glucose and methanol MOPS minimal medium (n=3, biological repeats) (c) Fermentation profile of SM1, SM6 and BW25113 strain. (n=3, biological repeats; error bars= S.D.)

There are several genes or promoters interrupted by insertion sequences, or frameshift mutations causing knockout or knockdown effects, including *rhlB*, *gtrS*, *quuD, yclG, gatY, rusA,* and *gltA*. To probe the effect of these mutations on growth, we complemented these inserted genes with a copy of intact gene on BAC in SM6n and tested their effects on growth rate. Results (Fig. 8a) showed that the complementation by BAC-borne *gltA* and *rhlB* genes significantly reduced the growth rate of SM6n. This result indicates that the *gltA* insertion by IS2 in the promoter region and *rhlB* frameshift were beneficial for growth. GltA is the first enzyme in the TCA cycle, which produces NADH. As methanol is already an electron-rich substrate, more so than glucose. A strong TCA cycle would increase the burden of respiration that regenerates NAD. In the SM6 strain the *gltA* promoter was interrupted by IS2, but it did not completely abolish the *gltA* expression. Proteome data suggested that GltA protein was still produced, albeit at about 10% level compared to the wild-type strain (Fig. 7c). Thus, it is possible that by properly knocking down *gltA*, the cell could balance NADH flux better. The same promoter mutation actually existed in the SM1 strain, however, re-introducing a wild-type *gltA* copy did not affect SM1 growth significantly^5^.

The *rhlB* gene encodes an ATP-dependent RNA helicase. In SM6, this gene contains a frameshift mutation and abolishes its function. It has been reported that overexpression of *rhlB* increased fitness and buffered deleterious mutations in *E. coli* mutator strains^18^. The inactivation of *rhlB* probably increased the effect of *mutS* mutation and accelerated evolution. However, its exact role in SM6 requires further investigation.

SM6 acquired the fast-growth ability in methanol minimal medium compared to SM1 (Fig. 6a). Interestingly, SM6 grows faster in methanol than in glucose minimal media (Fig. 8b), suggesting that this strain has already changed its preferred substrate from sugars to methanol. In aerobic methanol and glucose conditions and anaerobic glucose conditions, SM6 produced formate, acetate, and ethanol as by-products (Fig. 8c) similar to the wild-type strain in glucose. Note that the parental SM1 strain could not grow in glucose minimal medium anaerobically, but SM6 has recovered this ability. Neither SM1 nor SM6 could grow in methanol, possibly because of electron imbalance under fermentative conditions.

## Discussion

Methanol growth using the RuMP pathway requires only three enzymes (Mdh, Hps, and Phi) that are not present in wild-type *E. coli.* It may deceivingly appear quite simple to convert *E. coli* to a synthetic methylotroph by introducing these three enzymes. However, the challenge was more significant than expected. Many groups reported methanol-dependent growth of *E. coli* ^19–21^, which can assimilate methanol but still requires other carbon sources. The next step was the development of methanol auxotrophy that requires methanol to grow, even though other carbon sources were also needed^4,6^. A complete synthetic methylotroph took much longer to develop. We previously identified that the methanol metabolic intermediate, formaldehyde, could accumulate during the growth or stationary phases^5^. Formaldehyde is a potent chemical that causes DPC and kills the cell. Thus, cells with severe DPC is difficult to evolve. We solved this problem and evolved a first-generation synthetic methylotroph (SM1) that can grow with methanol as the sole carbon source. However, its growth rate is still far below that of native methylotrophs.

We hypothesize that the residual DPC in SM1 is responsible for the slow growth. To solve the DPC problem requires balancing formaldehyde production from methanol and formaldehyde consumption through the RuMP cycle. In this work, we used a BAC system that exhibits a dynamic CNV property through unknown sequences in the *ddp* operon. The presence of such sequence caused the whole BAC plasmid to generate tandem repeat concatemers with a wide range of copy numbers through a *recA*-dependent mechanism. In parallel to our effect in understanding the mechanistic details, we employed this system to solve the DPC problem in synthetic methylotrophic *E. coli*. The dynamic CNV is reminiscent of the copy number engineering technique^22^ but with a different mechanistic basis.

Indeed, using the ddp-BAC system, we were able to alleviate the DPC problem that became a bottleneck in the evolution. Introduction of ddp-BAC::RHTTP to *E. coli* allowed the cell to grow and evolve in high methanol concentration. The presence of multi-copy RHTTP allowed the cells to accumulate beneficial mutations. Once other alternative pathways to alleviate DPC have emerged, the cell lowered the copy number by inactivating *recA*. After we re-introduced an active copy of the *recA* gene through a multi-copy plasmid, the copy number increased immediately, but the *recA*-containing plasmid was purged after a few passes. The copy number returned to a low level. These results demonstrate the dynamic tuning of copy numbers to fit the particular genetic background. This feature allows the evolution process to bypass bottleneck, and eventually a fast-growing strain SM6 emerged.

The SM6 strain grows with a doubling time of 3.5 hr, and exhibits no lag phase in re-growth. The resolution of the DPC issue was attributed to the balancing of formaldehyde production and consumption enzyme activities. The preferred carbon source of SM6 has shifted from glucose to methanol.

These changes may be related to 6 mutated global regulatory genes in SM6, including the truncated *proQ* and *ptsP* inherited from SM1, along with newly acquired mutations in *cra, ompR, ihfA*, and *gadW* (Fig.5). The *cra* (Catabolite Repressor/Activator) gene, formerly also known as *fruR*, regulates key enzymes in central carbon metabolism pathways. This could affect the overall metabolic flux distribution in the cell. The EnvZ/OmpR pair is a two-component regulatory system that is responsible for osmotic response, as it regulates the expression of outer membrane proteins (OMPs) like OmpF and OmpC. The mutation may have affected altered the osmotic pressure and membrane permeability to adapt to the uptake of methanol and formaldehyde toxicity.

One potential limitation of SM6 is that it cannot grow fermentatively in methanol minimal medium. This phenotype may be attributed to the electron-richness of methanol. It is difficult to balance the electron with typical fermentation products. Further work is in order to overcome this problem effectively.

## Acknowledgements

We thank the following core facilities at Academia Sinica: the NGS High Throughput Genomics Core Facility at the Biodiversity Research Center, the Imaging Core Facility at the Institute of Molecular Biology Center, Bio-Imaging Core Facility at the Institute of Biological Chemistry, and the Academia Sinica Common Mass Spectrometry Facilities for Proteomics and Protein Modification Analysis at the Institute of Biological Chemistry. The NGS Core Facility and mass spectrometry Facility are supported by the Academia Sinica Core Facility and Innovative Instrument Project, Taiwan (AS-CFII-108-114 and AS-CFII-108-107)

## Author Contribution

LYN, FYC and JCL conceived the project. LYN developed the ddp-BAC system. FYC, LYN, HWJ, KYS conducted the evolution process. KYS analyzed the ddp-BAC nanopore sequencing results. FYC analyzed the NGS and proteomics data. LYN analyzed the formaldehyde concentration of the SM strains. FYC and HWJ worked on ddPCR. FYC conducted the TEM DPC experiment. LYN and FYC worked on characterizing the mutations of the SM6 strain. FYC worked on GC-MS analysis metabolites. HWJ worked on the library prep for all DNA sequencing samples. All authors worked on designing constructs, cloning, or participated in writing and preparation of the manuscript and figures.

## Materials and Methods

### Plasmid construction

KODone polymerase (TOYOBO) was utilized for all PCR reactions. The NEBuilder 2x HiFi DNA Assembly Master Mix (NEB) was employed for Gibson assembly to construct plasmids. Subsequently, the plasmids were introduced into DH5α cells (NEB) cultivated in lysogeny broth (LB medium) or Terrific Broth (TB medium) supplemented with appropriate selective antibiotics. The plasmid sequences were verified by Sanger sequencing or nanopore sequencing.

### Strain construction

For genome editing of methylotrophic strain, we used Tn7-like transposons-mediated CRISPR system from Vo, Phuc Leo H., et al. ^23^. All plasmids transformed into the host cells were done by electroporation (Biorad).

### Media for methylotrophic growth

MOPS EZ buffer (Teknova) was used as a MOPS minimal medium, which contained 40 mM MOPS, 3.02 nM CoCl_2_, 0.962 nM CuSO_4_, 50 mM NaCl, 9.5 mM NH_4_Cl, 0.525 mM MgCl_2_, 4 mM tricine, 1.32 mM K_2_PO_4_, 0.276 mM K_2_SO_4_, 0.01 mM FeSO_4_, 0.5 mM CaCl_2_, 40 nM H_3_BO_3_, 8.08 nM MnCl_2_, 0.974 nM ZnSO_4_, and 0.292 nM (NH_4_)_2_MoO_4_. MOPS-based methanol minimal medium was composed of methanol, 50 mg/mL chloramphenicol, 1 mM IPTG and MOPS EZ buffer. Chloramphenicol was dissolved in methanol. A vitamin mix is prepared where the following final concentration is reached in the medium: 8.19 μM biotin, 10.49 μM calcium pantothenate, 4.53 μM folic acid, 40.94 μM nicotinamide, 13.29 μM riboflavin, 14.82 μM thiamine hydrochloride, and 0.07 μM vitamin B12.

### Digital droplet PCR (ddPCR) to analyze copy number

Genomic DNA from *E. coli* was extracted either manually using the Puregene Kit from Qiagen or automatically purified using the magnetic bead system from Taco™ mini (GeneReach company, TW). The concentration of genomic DNA (gDNA) was quantified using the BioTek Synergy H1 Plate Reader (Agilent Technologies), aiming for a concentration of 500 ng. The gDNA was fragmented by incubating with the restriction enzymes PstI/HindIII at 37°C for 15 minutes in a total volume of 50 μl. The concentration of DNA fragments was diluted to 25 pg/μl, and quantification was performed using the Qubit™ 1X dsDNA High Sensitivity assay kit (Thermo Fisher Scientific).

A ddPCR solution was prepared containing ddPCR master mix and probes and primers for the target gene labeled with FAM, reference gene 1 labeled with HEX, and reference gene 2 labeled with Cy5. ddPCR was performed using the Naica® system for Crystal Digital PCR™ (Stilla Technologies). 10-25 pg of DNA was mixed with the ddPCR solution and injected into either a Sapphire chip for 12 samples or a Ruby chip for 48 samples. Partitioning and PCR were carried out using the Geode device, followed by chip reading using the Prism 6 device. Data analysis was conducted using the Crystal Miner data analysis software. Copy number was calculated by dividing the absolute concentration of the target gene by the absolute concentration of the reference gene.

### Next-generation sequencing and Nanopore sequencing

DNA samples for sequencing are extracted either using the Nanobind CBB kit (PacBio, USA)], the Puregene kit (Qiagen, Germany), or automatically extracted using the QIACUBE DNeasy (Qiagen, Germany). For NGS, 200ng of DNA is used for library preparation with the Illumina DNA Prep Kit (Illumina, USA). The quality of DNA libraries is assessed using the 5200 Fragment Analyzer and sequenced on the Illumina MiniSeq or NextSeq with a read length of 150bp paired-end. For normal reads (N50 about 7kb) nanopore sequencing, 400ng of DNA is used for library preparation with the Rapid Barcoding Kit 96 (Nanopore, UK). For long reads (N50 about 15kb) nanopore sequencing, 1ug of DNA is used for library preparation with the Native Barcoding Kit 24. Sequencing is performed using a MinION Flow Cell (R9.4.1 or R10.4.1) and MinION MK1B, with real-time basecalling carried out using the MinKNOW software on a DELL workstation computer equipped with a high-performance GPU RTX 3090. All SNP calling and variant detection, annotations were all performed by Geneious Prime software.

### Microscopy and live cell imaging

The real-time imaging of *E. coli* was conducted by IX83 Inverted Microscope (IX83, Olympus, Tokyo, Japan) equipped with an Olympus UPlanXApo 60x / 1.42 Oil Ph3, Infinity / 0.17 Microscope Objective, and an ORCA-Flash 4.0 V3 Digital CMOS camera (Hamamatsu Photonics). Following cultivation of *E. coli* to OD600 to 0.1, a 10 μL aliquot was applied onto a µ-Dish 35 mm, high Glass Bottom (Cat. No: 81158, ibidi) loaded with a 1% (w/v) agarose base, and subsequently positioned within the microscope setup for imaging at 37°C for 4 hours. Cell image processing was conducted using Fiji software.

### Flow cytometry analysis

Flow cytometry analysis was performed using a CytoFLEX S flow cytometer (Beckman), equipped with a 488 nm blue laser emitting at 51 mW, which facilitated the detection of forward scatter (FSC), side scatter (SSC), and fluorescence signals. All the samples were taken from bacterial cultures grown in LB overnight, and then were appropriately diluted in autoclaved 1x phosphate-buffered saline (PBS) and adjusted to a concentration of approximately 1.5 x 10^6^ cells/mL. To minimize background noise, SSC thresholds for all collected samples were set at a value of 1000. Data acquisition was conducted at a rate exceeding 1000 events per second (approximately 30 µL/s), with an abort rate of less than 10%. Fluorescence distribution histograms were generated using FCS Express software (De Novo Software).

### Growth assay and laboratory evolution of synthetic methylotrophic strain

Methylotrophic strains were evolved in 3 mL tubes at 37°C and 250 rpm using a New Brunswick Scientific Innova 44 shaker. These strains were then transferred to fresh medium at an initial OD ranging from 0.05 to 0.2 once they reached the stationary phase. OD values were measured using a GENESYS™ 30 Visible Spectrophotometer. To optimize growth conditions, we employed the RTS-8 Plus system (Biosan) to monitor growth rates under high-aeration conditions.

### The detection of formaldehyde in culture medium

50 µL of bacterial culture was filtered through an Amicon Ultra-0.5 centrifugal filter(50 kDa, Merck), and then incubated with 50 µL of 0.1% 2,4-Dinitrophenylhydrazine (DNPH) and 2 µL of 85.4% H_3_PO_4_ at room temperature for 10 minutes. After incubation, 2 μL of the sample was injected into the HPLC system for separation by a Triart C18 ExRS column (1.9 μm, 2.1 × 100 mm, YMC) with a constant flow rate of 0.3 mL/min at 50 °C. Eluent A consisted of 50% acetonitrile in water (50 v/v) with 0.4 mM ammonium fluoride, while eluent B contained 100% methanol with 0.4 mM ammonium fluoride. The HPLC program was set as follows: 100% A (0.0–3.5 min), 0 to 98% B (3.5–5.5 min), 98% B (5.5–10.0 min), 98 to 0% B (10.0–10.5 min). A heated ESI probe was equipped for ionization and operated in negative ion mode. The data were processed and analyzed using LabSolutions software (Shimadzu).

### Transmission Electron Microscopy (TEM) analysis to characterize DPC

Purified DPC complexes, approximately 500 ng of DNA, were mounted onto activated 300-mesh copper grids coated with carbon-stabilized formvar (Ted Pella). This mounting process was operated at room temperature for a duration of 1 minute. Following the removal of excess liquid with filter paper blotting, the samples were stained with a 2.5% uranyl acetate solution for 1 minute and air-dried naturally at room temperature. TEM images were captured using a Talos L120C. The magnification ratio was set to range from 2700x to 15000x for optimal visualization.

### Enzyme Assay

Enzymes from the RuMP pathway (including Hps, Phi, Tkt, Tal, Rpe, Rpi) were purified from a lac-induced His-tagged overexpression system in *E. coli.* The cells were inoculated in 5ml LB and then grown over night at 37°C. On the following day, the overnight culture was inoculated into a 100ml LB at 1% and then grown at 37°C. 0.1mM IPTG was then added when the culture reached OD 0.4∼0.6 at log phase. The culture was then grown at 20°C 250rpm overnight, and then collected by centrifugation. The cells were then resuspended with 10mM Phosphate buffer solution, pH 7.4, and broken by a Sonicator (Q600) on ice. After 1hr of 15000rpm centrifugation, the protein solution was then filtered by a 0.22 μM filter, and then purified by a Biorad NGC using a IMAC column (Roche). The crude extracts are prepared as followed. 2ml of normalized OD 1 culture were harvested from SM6 and SM1 when grown at OD1 and OD3 respectively, where dilution is done by pure PBS. The cultures are then centrifuged at 7000rpm, where supernatant is discarded, and the pellets are resuspended in 10mM Phosphate Buffer, pH 7.4 at 700ul. The mixture was then sonicated, and the supernatants (crude extracts) were collected by 1 hour of centrifugation at 18000rpm. All samples are normalized to a concentration of 1mg/ml. The Pgi and Zwf enzymes were obtained commercially from Roche. For Hps/ Phi Assay, the following enzymes were utilized: Hps, Phi, Rpi, Pgi, Zwf with a NADPH readout with 100mM Phosphate buffer (pH7.4), 10mM formaldehyde, 4mM R5P, 1mM thiamine pyrophosphate, 2mM MgCl2. For Tkt/Tal/Rpi Assay, the following enzymes were utilized: Tkt, Tal, Rpe, Rpi, Pgi, Zwf with a NADPH readout with 100mM Phosphate buffer (pH7.4), 4mM R5P, 1mM thiamine pyrophosphate, 2mM MgCl_2_. Mdh is tested by a Nash reaction assay, similar to our previous report ^24^.

### Proteomics analysis

Samples were processed by homogenizing for 2 minutes on ice using a UP50H homogenizer with a sonotrode 3 (Hielscher, Ultrasound Technology). Following homogenization, samples were centrifuged at 17,000X g for 20 minutes at 4 °C for clarification. The supernatant was then collected, and protein content was quantified using absorbance of 280nm and Bradford assay. For mass spectrometry analysis, proteins (50 µg from each sample) were first reduced with 10 mM dithiothreitol (DTT) (Sigma-Aldrich, St. Louis, MO, USA) and subsequently alkylated using 50 mM iodoacetamide (Sigma-Aldrich, St. Louis, MO, USA). Proteins were then digested at room temperature for 3 hours with Lys-C endopeptidase (Wako, Tokyo, Japan) in a ratio of 1:50 by weight, followed by an overnight digestion with trypsin (sequencing grade, Promega Corporation, Madison, WI, USA) at a 1:100 weight ratio at 37 °C. Digested samples and extract detergent were acidified by adding trifluoroacetic acid (TFA) and ethyl acetate (Wako), respectively. Peptides were then desalted using SDB-XC Empore disk membranes in StageTips (GL Sciences, Tokyo, Japan) and measured with the Pierce™ Quantitative Colorimetric Peptide Assay (Thermo Fisher Scientific). Equal quantities of purified peptides from each sample were then dried and prepared for TMT labeling. Peptides were reconstituted in 200 mM HEPES and treated with TMT 18-plex™ Isobaric Label Reagent Set (Thermo) at room temperature for 1 hour. The reaction was quenched using 0.33% hydroxylamine for 15 minutes at room temperature. The labeled samples were then pooled, dried, and fractionated on basic reversed-phase SDB-XC StageTips. The resultant peptide fractions were further purified using Pierce™ C18 Spin Tips (Thermo Scientific). The samples were dried and redissolved in 2% acetonitrile (ACN) and 0.1% formic acid. Peptide analysis was conducted using an Orbitrap Fusion Lumos Tribrid mass spectrometer (quadrupole-ion trap-Orbitrap, Thermo Fisher Scientific, San Jose, CA, USA), coupled with an Ultimate 3000 nanoLC system (Thermo Fisher Scientific, Bremen, Germany). Peptides were loaded onto a C18 Acclaim PepMap NanoLC column (25 cm length, 75 µm inner diameter, Thermo Fisher Scientific, San Jose, CA, USA) packed with 2 µm particles and a 100 Å pore size. The mobile phase A consisted of 0.1% formic acid in water, and the mobile phase B was 99.9% ACN with 0.1% formic acid. Gradient elution was performed over 50 min, increasing ACN from 2% to 40% at a flow rate of 300 nL/min. The mass spectrometer was operated in data-dependent mode, automatically alternating between MS1 and MS/MS acquisitions. MS1 spectra were recorded in the Orbitrap at a resolution of 120,000 for m/z segments from 350 to 1700, with an automatic gain control (AGC) target of 5 × 10^5^ and a maximum injection time of 50 ms. The instrument was configured for top-speed mode with 3-second cycles for both survey and MS/MS scans. Peptide ions ranging from charge states 2 to 7 were isolated with a width of 1.4 Da for higher-energy collisional dissociation (HCD) fragmentation at a normalized collision energy (NCE) of 38%. Fragment spectra were collected in the Orbitrap analyzer at a resolution of 60,000. An AGC target of 5 × 10^4^ was set for MS/MS analyses, with dynamically excluded previously selected ions for 60 s. The proteomics data were processed and analyzed by Proteome Discovery (PD) and Perseus software. Enriched biological functions and processes of identified differential expressed proteins were performed by Gene Ontology (GO) analysis.

### The measurement of fermentation products

Ethanol, formate and acetate were analyzed using an Agilent 7890B Gas chromatography with a 5977 mass spectrophotometer GC-MS with a SH-I-5MS column (Shimadzu). Helium is used as the gas carrier with constant pressure at 7.0633 psi. The thermal cycle is as follows: initial temperature: 40 °C, 2 minutes, ramp rate 10 °C /min to 60 °C, 40 °C /min to 220 °C, 2 minute final hold.

### Rifampicin-based mutation assay

1 mL of bacterial cultures were harvested and sprayed on rifampicin plates at the concentration of 25 µg/mL. Meanwhile, the strain was seeded on LB plates to normalize cell number. The mutation rate was determined by calculating the colony-forming units per mL on rifampicin plates, normalized with CFU/mL on LB plates.

### Analysis of ^13^C labelling pattern of amino acids and intracellular primary metabolites from methanol-growing SM1

Direct analysis of amino acids was conducted by a UHPLC-MS system with Agilent 1290 Infinity II ultra-high performance liquid chromatography (UHPLC) system coupled online to Agilent 6545 quadrupole time-of-flight (Q-TOF) mass spectrometer (Agilent Technologies). Samples were retained by using ACQUITY UPLC BEH amide column (1.7 μm, 2.1 × 100 mm, Waters), with a constant flow rate of 0.3 mL/min and the injection volume of 2 μL. The mobile phases were composed of H2O (eluent A) and 90% acetonitrile (eluent B) respectively, where both eluents contained 15 mM ammonium acetate and 0.3% ammonium hydroxide. The Dual Agilent Jet Stream (AJS) electrospray ionization (ESI) was selected as the ion source, and the instrument was operated in positive and negative full-scan mode, collecting m/z 60–1500. After-run analysis was processed using Agilent Qualitative Analysis 10.0 and Agilent Profinder 10.0 software (Agilent Technologies), which included chromatogram acquisition, detection of mass spectral peaks, and waveform processing. In another method, amino acids were derivatized by o-phthalaldehyde (OPA, Sigma), 3-mercaptopropionic acid (MPA, Sigma) and 9-fluorenylmethyl chloroformate (FMOC, Sigma), and then analyzed by LC-MS/MS. Specifically, 30 μL of MPA and 15 μL of OPA were mixed with 5 μL appropriately diluted sample and placed at room temperature for 1 minute. After primary amine derivatization, the sample were further reacted with FMOC for 2 minutes. The LC-MS/MS system is based on Shimadzu LCMS-8045 triple quadrupole mass spectrometer equipped with high performance liquid chromatography system (HPLC, Nexera X2 LC-30AD, Shimadzu). 2 μL of sample was injected into HPLC system, and the derivatized amino acids were separated by Triart C18 ExRS column (1.9 μm, 2.1 × 100 mm, YMC) with a constant flow rate at 0.3 mL/min, 50 °C. The eluent A consisted of 2.38 mM tributylamine, 1.28 mM acetate and 0.4 mM ammonium fluoride, while the eluent B contained 100% methanol with 0.4 mM ammonium fluoride. The HPLC program is as follows: 5-25% B (0.0–4.0 min), 25–98% B (4.0–10.0 min), 98% B (10.0–20.0 min), 98–5% B (20.0-20.2 min) and 5% B (20.2–25.0 min). Heated ESI probe was equipped for ionization and was operated in negative ion mode. The data were processed and analyzed by LabSolutions software (Shimadzu). Intracellular primary metabolites were analyzed by Shimadzu LCMS-8045 triple quadrupole mass spectrometer. 2 μL of sample was separated by a 15 cm Triart C18 ExRS column (1.9 μm, 2.1 × 150 mm, YMC) in HPLC system with a constant flow rate at 0.3 mL/min 50 °C. The composition of mobile phase, HPLC program, ionization mode and data processing were the same as the method for analyzing derivatized amino acids.

## Supplementary Figures and Tables

**Supplementary Figure 1.**
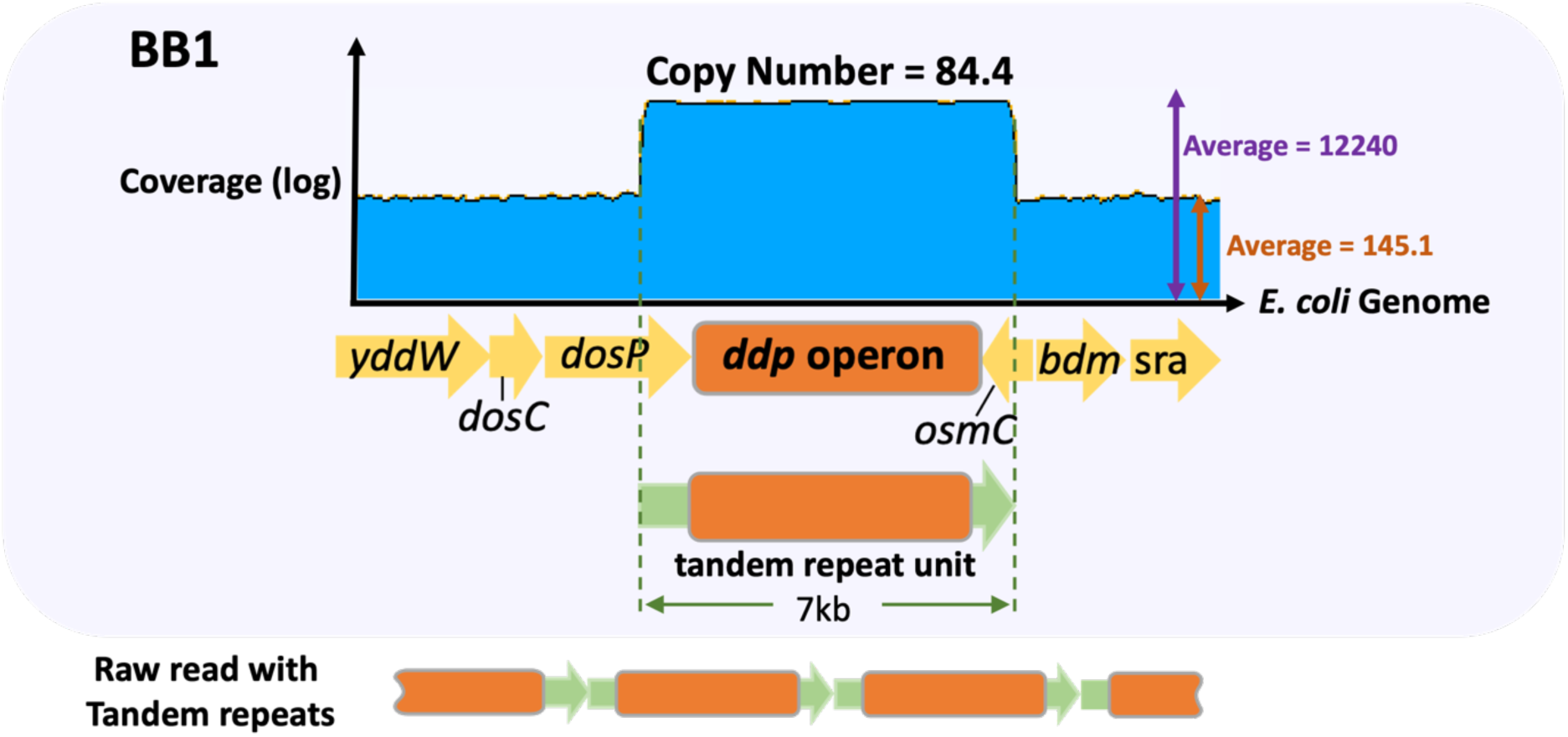
The BB1 strain contains a 7kb region with an elevated copy number (> 80). This strain spontaneously emerged and was co-evolved with the SM1 strain during the methanol evolution process in our previous work (Chen et al, Cell 2020). Nanopore sequencing revealed that this region consists of tandem repeats with fixed boundaries.

**Supplementary Figure 2.**
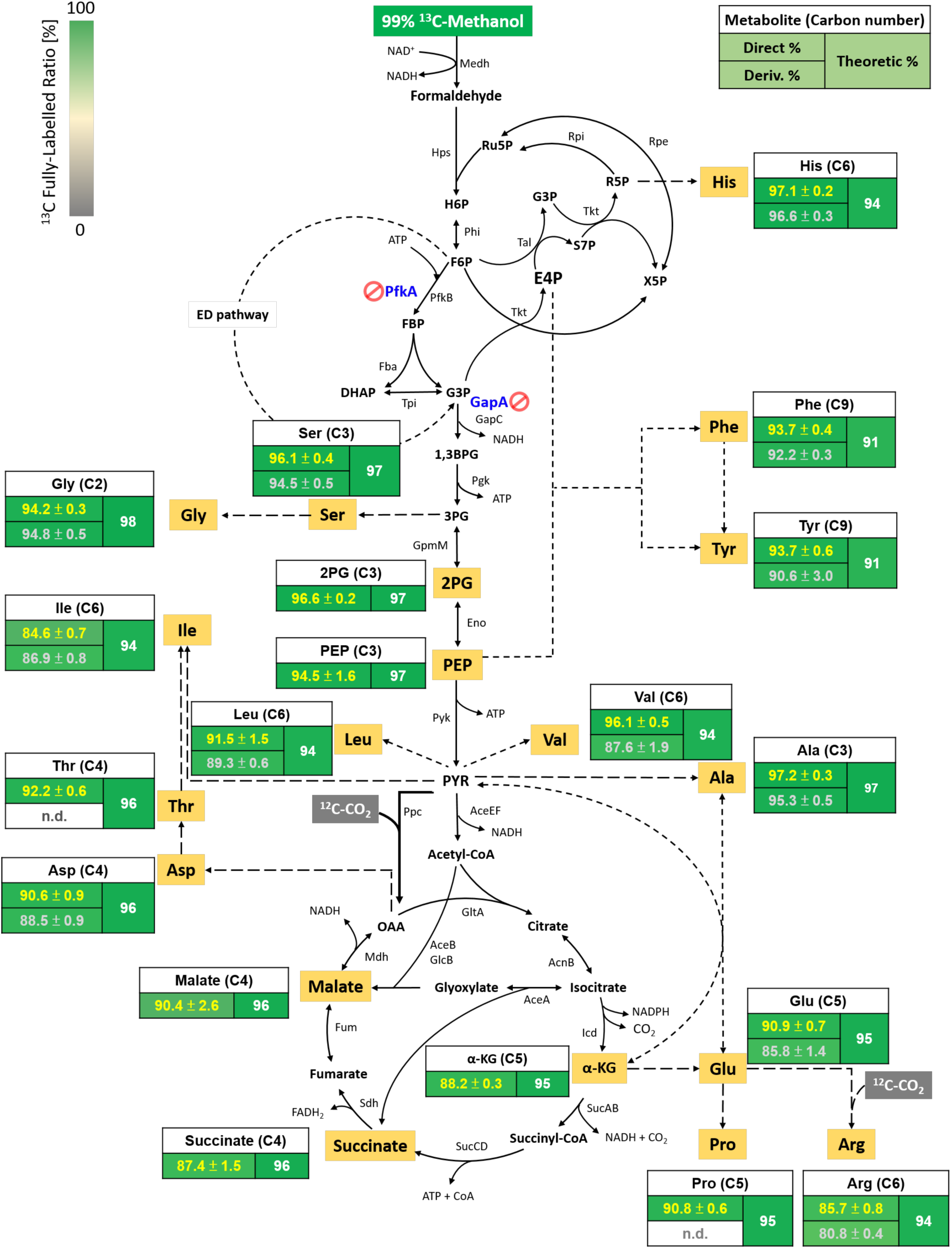
^13^C labelling pattern of biomass hydrolysates and intracellular primary metabolites in *E. coli* SM1 growing on MOPS based minimal medium with 13C-methanol as the sole carbon source. “Direct” refers to biomass amino acid labelling results directly measured by LC-qTOF, or intracellular primary metabolites directly measured by LC-MS/MS. “Deriv.” refers to amino acid measured by LC-MS/MS after derivatization by OPA/MPA/FMOC. “Theoretic” refers to the theoretic maximum labelling percentage calculated by the purity of the acquired 99% ^13^C methanol and the natural isotope abundance of other atoms. (n=3; data present mean ± S.D.; n.d = not detected)

**Supplementary Figure 3.**
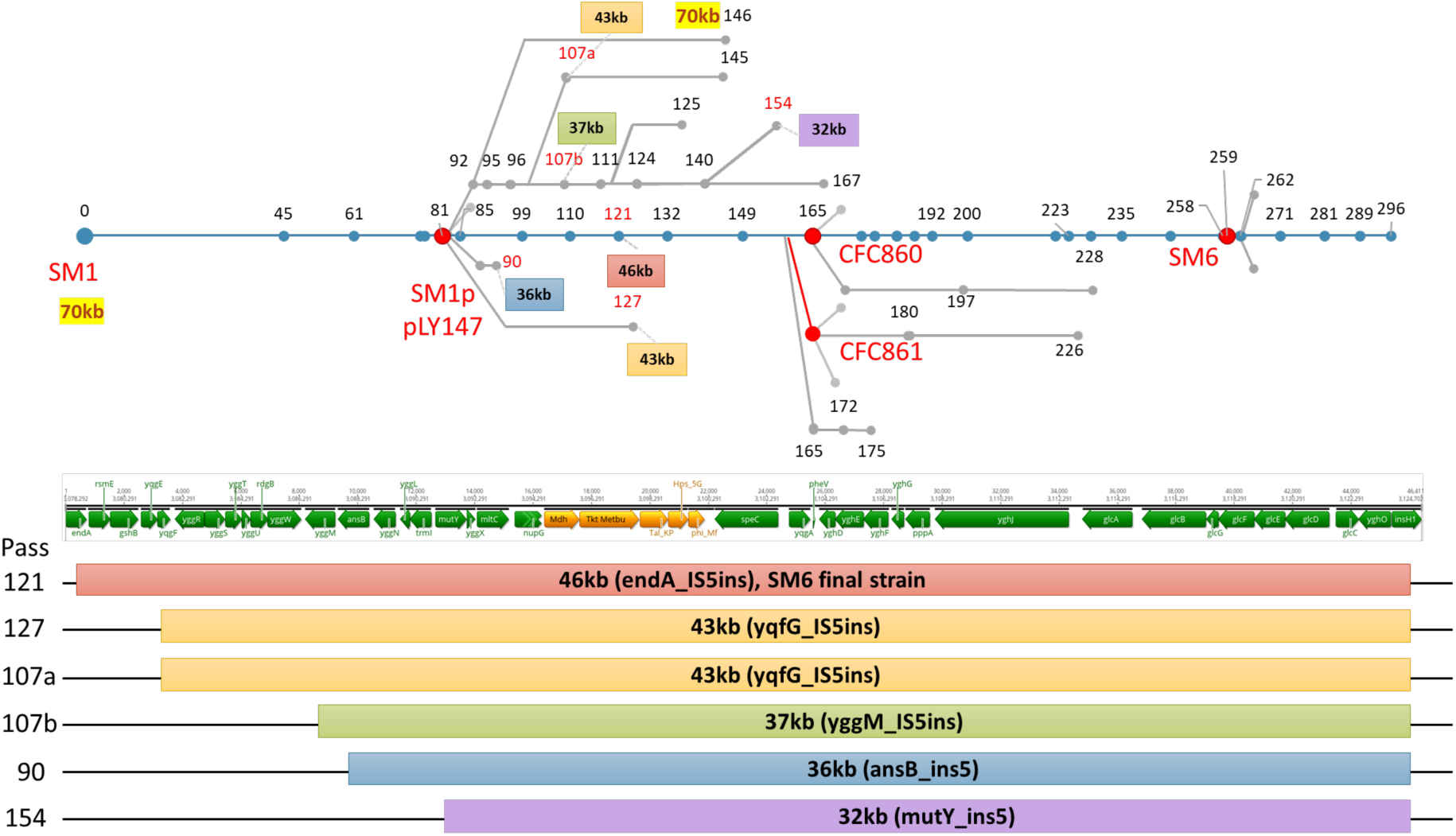
Unit size distribution of the IS5-mediated tandem repeats from the original 70kb tandem repeat of SM1. The colored regions correspond to the same-color-coded pass numbers shown in the evolution lineage graph.

**Table S1.**
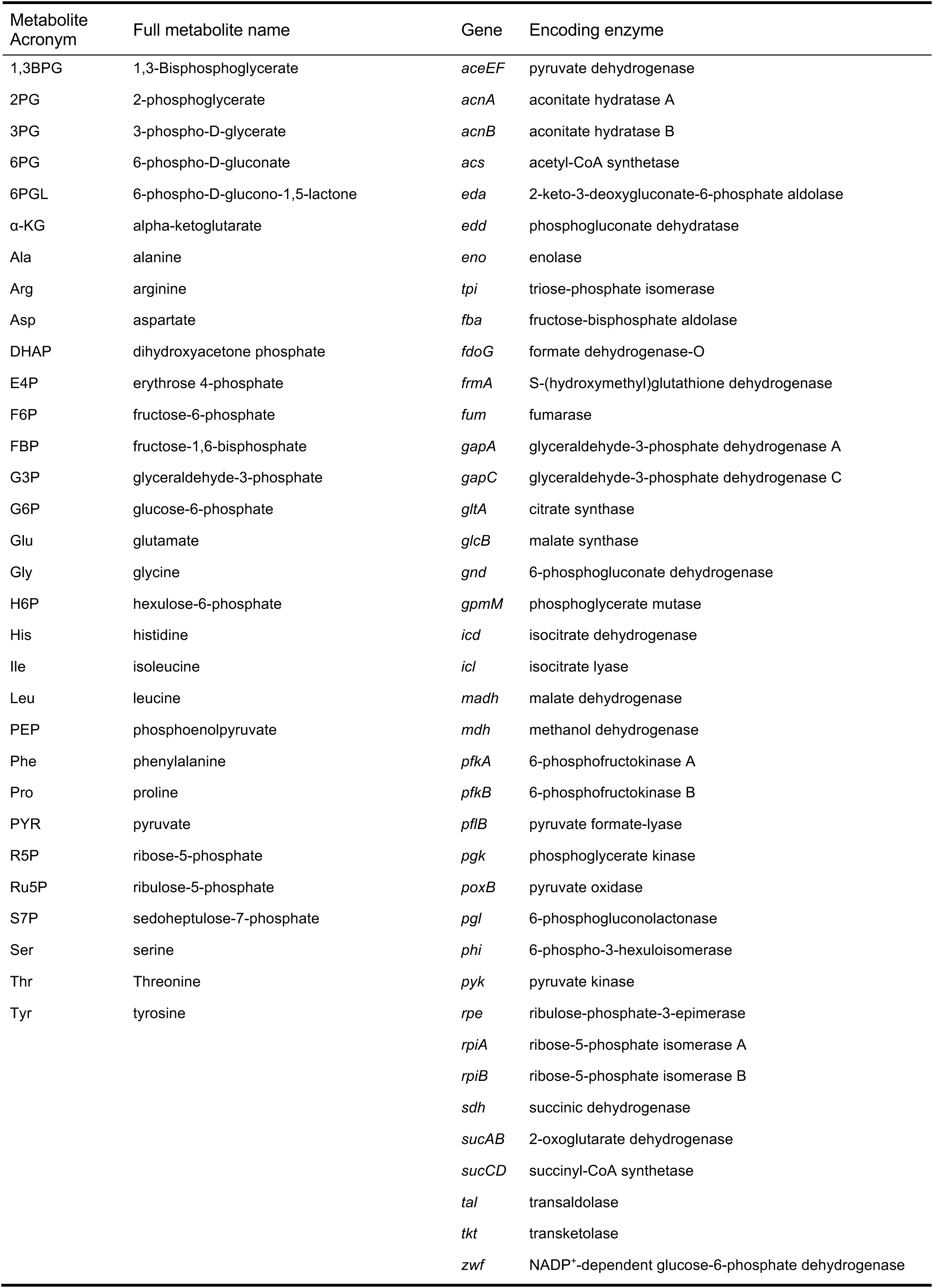
Metabolite and gene list.

**Table S2.** Genotype of strains and cultures.

|  | Codon Change | Non-coding Region | Indel Codon Change | Large Genome change | IS-mediated tandem repeat |
| --- | --- | --- | --- | --- | --- |
| SM1 | *araG* (I275R)<br>*rpoA* (G315C)<br>*xylR* (K320Q)<br>*rpoC* (S733F)<br>*proQ* (E12*)<br>*icd* (D398E)<br>*icd* (D410E) | *gltA*_IS2<br>*ptsH*_IS2 | *relE*_IS2<br>*yhcM*_IS2<br>*smf*_IS2<br>*elfG*_IS3 | *ΔrpiA*<br>*ΔrpiB* |  |
|  |  |  | *fdoG*<br>(2,194_del G)<br>[Frame shift] | 2,093,044-2,095,229<br>*ugd* N-terminal and operon *gnd*, *wbbL* deletion | 130k<br>(*rrsA* to *rrlB*) duplicate |
|  |  |  | *frmA*<br>(383_ins CCCG)<br>[Frame shift] | *stfP*_291-*stfE*_3 transversion<br>*ΔgapA::gapC*<br>*ΔpfkA* | 240k<br>(*yqiG* to *smf*) duplicate |
|  |  |  | *ugd*<br>(1-51_del)<br>[Lost start codon] | 1,191,676-1,206,868<br>cryptic prophage *e14* deletion | 70k<br>(*yggE* to *yghO*) 5 fold |
|  |  |  | *pgi*<br>(705_del GTTGCAAAACAC)<br>[Codon:236-239_delVAKH] |  |  |
| SM1p | *Cra* (E190Q)<br>*ihfA* (V78E) |  | *kdsC*_IS2<br>*tauB*_IS2<br>*yaaA*_IS2<br>*yiaA*_IS2 |  | 130K disappeared<br>240K disappeared |
| CFC860 | *bglX* (W430G)<br>*cra* (E190Q)<br>*ihfA* (V78E)<br>*queE* (D180Y) |  | *kdsC*_IS2<br>*rimJ*_IS2<br>*tauB*_IS2<br>*yaaA*_IS2<br>*yiaA*_IS2<br>*yqeI*_IS2<br>*endA*_IS5 | *ddpA* on both genome and pLY147 was inserted with pJ23102: *mdh* |  |
| SM6 | *acrB* (A303S)<br>*bglX* (W430G)<br>*cra* (E190Q)<br>*dapF* (A211V)<br>*ebgA* (V339I)<br>*fliE* (P28S)<br>*gadC* (A28T)<br>*glmU* (N283S)<br>*hemY* (G335D)<br>*ihfA* (V78E)<br>*nuoC* (T545I)<br>*queE* (D180Y)<br>*rpoB* (V2F)<br>*rpoD* (F401S)<br>*yhbU* (A89V) |  | *cbrB*_IS2<br>*gtrS*_IS2<br>*kdsC*_IS2<br>*mutS*_IS2<br>*rimJ*_IS2<br>*tauB*_IS2<br>*yaaA*_IS2<br>*yiaA*_IS2<br>*ylcG*_IS2<br>*yqeI*_IS2<br>*quuD*_IS3<br>*endA*_IS5 |  |  |
|  |  |  | *gadW*<br>(86 ins G)<br>[Frame shift] |  |  |
|  |  |  | *rhlB*<br>(25 del T)<br>[Frame shift] |  |  |
|  |  |  | *ygiQ*<br>(714 ins T)<br>[Frame shift] |  |  |

**Table S3.**
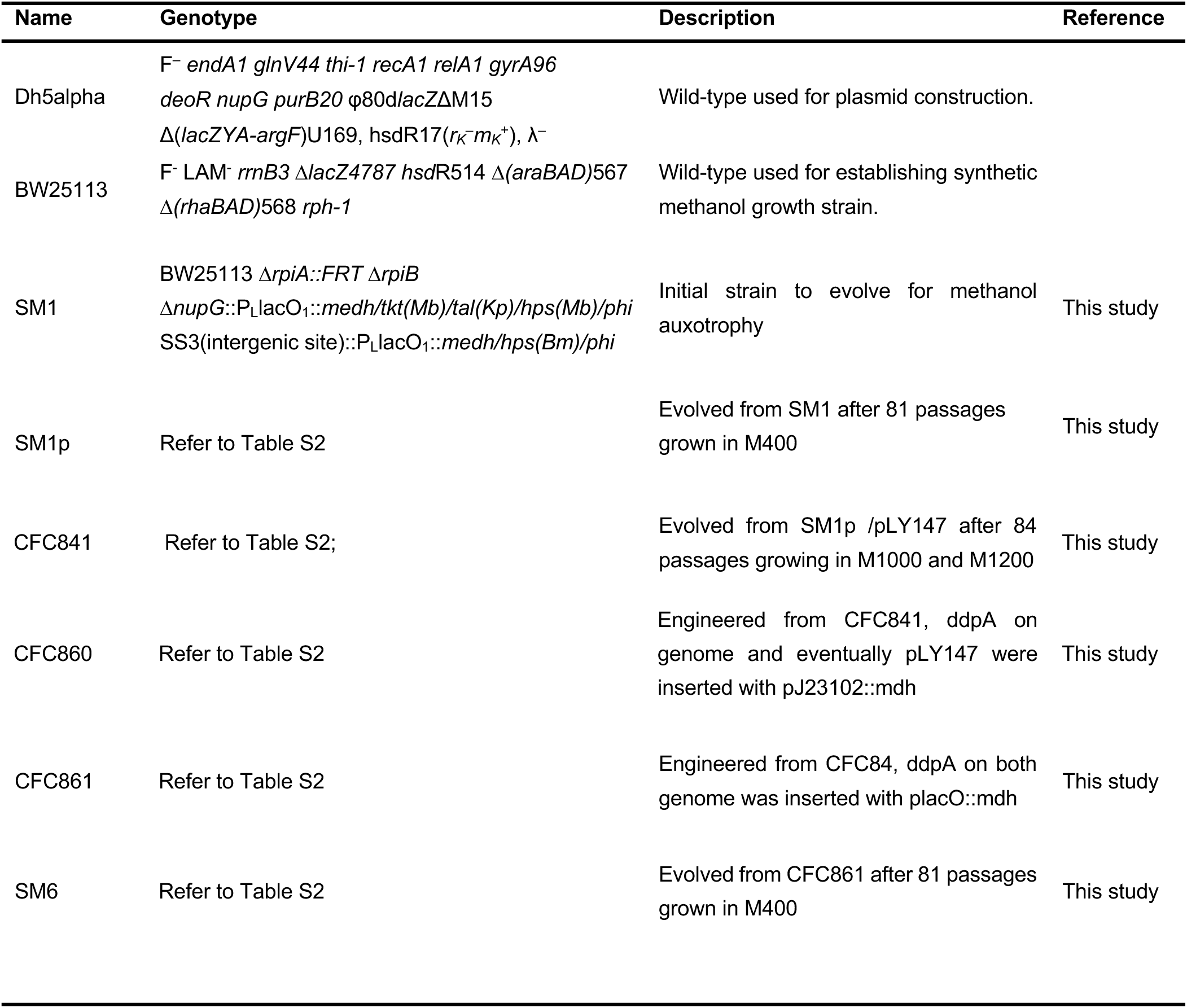
Strain list.

